# Mast cell-specific receptor/corticotropin-releasing factor axis regulates alcohol withdrawal-associated headache

**DOI:** 10.1101/2021.05.21.445199

**Authors:** Hyeonwi Son, Yan Zhang, John Shannonhouse, Hirotake Ishida, Ruben Gomez, Armen Akopian, Yu Shin Kim

## Abstract

Rehabilitation from alcohol addiction or abuse is challenging due to alcohol withdrawal symptoms. Headache is a severe alcohol withdrawal symptom that frequently contributes to rehabilitation failure. Despite the need for treating alcohol withdrawal-induced headache, there is no appropriate therapeutic option available. Development of improved therapeutics will depend on obtaining a clearer understanding of alcohol withdrawal-induced headache pain mechanisms. Here, we show that the mast cell-specific receptor MrgprB2 controls development of alcohol withdrawal-induced headache. Withdrawing alcohol from alcohol-acclimated mice induces strong headache behaviors, including facial allodynia, facial pain expressions, and reduced walking movement, symptoms often observed in humans suffering from headache. Observed pain behaviors were abolished in MrgprB2-deficient mice. We observed *in vivo* spontaneous activation and hypersensitization of trigeminal ganglia neurons in alcohol withdrawal mice, but not in MrgprB2-deficient mice. Corticotropin-releasing factor (CRF) was increased in dura mater after alcohol withdrawal. Injection of CRF into dura mater resulted in activation of trigeminal ganglia neurons and vasodilation, which was accompanied by headache behavior. In cells, CRF evoked Ca^2+^ transients via MrgprB2 or human MrgprX2. The results indicate that alcohol withdrawal causes headache via mast cell degranulation in dura mater. The process is under control of MrgprB2/MrgprX2, which would appear to represent a potential target for treating alcohol withdrawal-related headache.

## Introduction

Alcohol is an addictive substance, and approximately 380 million people suffer from alcohol abuse or dependence (WHO, 2016). Global disasters, such as terrorism, economic adversity, and the COVID-19 respiratory epidemic, are associated with increased alcohol consumption and increased vulnerability to development of risky drinking behaviors^1^. Rehabilitation from alcoholism is of critical importance to managing alcohol dependency of a large portion of the population. However, the rehabilitation process is hindered by alcohol withdrawal symptoms, specifically, headache^2–4^. The temporary relief from alcohol withdrawal-induced headache pain derived from resumption of alcohol consumption is a driving force failure to break the addiction cycle^5,6^, seriously affecting quality of life and aggravating alcohol dependence. Despite a major unmet medical need for treating alcohol withdrawal-induced headaches, there is no appropriate therapeutic option available. To develop better therapeutics, it will be necessary to obtain a clearer understanding of alcohol withdrawal-induced headache pain mechanisms.

Headache is initiated from the activation of trigeminal ganglia (TG) neuronal afferents and vasodilation in dura mater. Processes that cause local inflammation of dura mater sensitize peripheral afferents of TG neurons^7–9^, and their activation by mechanical and chemical stimuli contribute to the progression of general headaches^7,10^. Mast cell degranulation in dura mater has been implicated in local inflammation and nociceptive afferent activation of TG neurons and vasodilation^11,12^, suggesting that activated mast cells in dura mater may mediate headache. Mast cells are located proximal to peripheral nerve endings and peripheral blood vessels in dura mater, where they can be activated by various secretagogues to release proinflammatory cytokines^13,14^. Mas-related G-protein-coupled receptor B2 (MrgprB2), which is selectively expressed on connective tissue mast cells, is activated by basic secretagogues^15^, and mediates neurogenic inflammation pain^16–18^. Chronic alcohol consumption induces increased mast cell numbers and degranulation in peripheral tissues, where mast cells could mediate inflammation ^19–21^. In view of these observations, we tested whether mast cell-specific MrgprB2 contributes to alcohol withdrawal-induced headache.

## Results

### Mast cell receptor MrgprB2 mediates alcohol withdrawal-induced headache behaviors

In a two-bottle (10% ethanol vs. water) voluntary choice paradigm, mice exhibited a preference for ethanol (Extended Data Fig. 1a). Neither food intake (Extended Data Fig. 1c) nor body weight (Extended Data Fig. 1d) were affected. Ethanol preference increased gradually over the three week study (Extended Data Fig. 1b), indicative of developing ethanol dependence. Withdrawal from voluntary ethanol access after 3 or 8 weeks induced mechanical hypersensitivity in the periorbital area, which was maintained for up to 4 days (Fig. 1a and Extended Data Fig. 3o). Moreover, the grimace pain score was significantly increased 24 hours after alcohol withdrawal (Fig. 1b and Extended Data Fig. 1e). Reduced exploratory behaviors in the open field test after alcohol withdrawal in mice (Fig. 1c and Extended Data Fig. 1f) correlated with avoidance of physical activity described in migraineurs^22^. However, the headache behaviors induced by alcohol withdrawal were abolished in MrgprB2-deficient (MrgprB2 KO) mice (Fig. 1a-c). Thus, withdrawal from ethanol resulted in headache behaviors, and indicated that mast cell-specific MrgprB2 likely mediates alcohol withdrawal-induced headache.

**Fig. 1.**
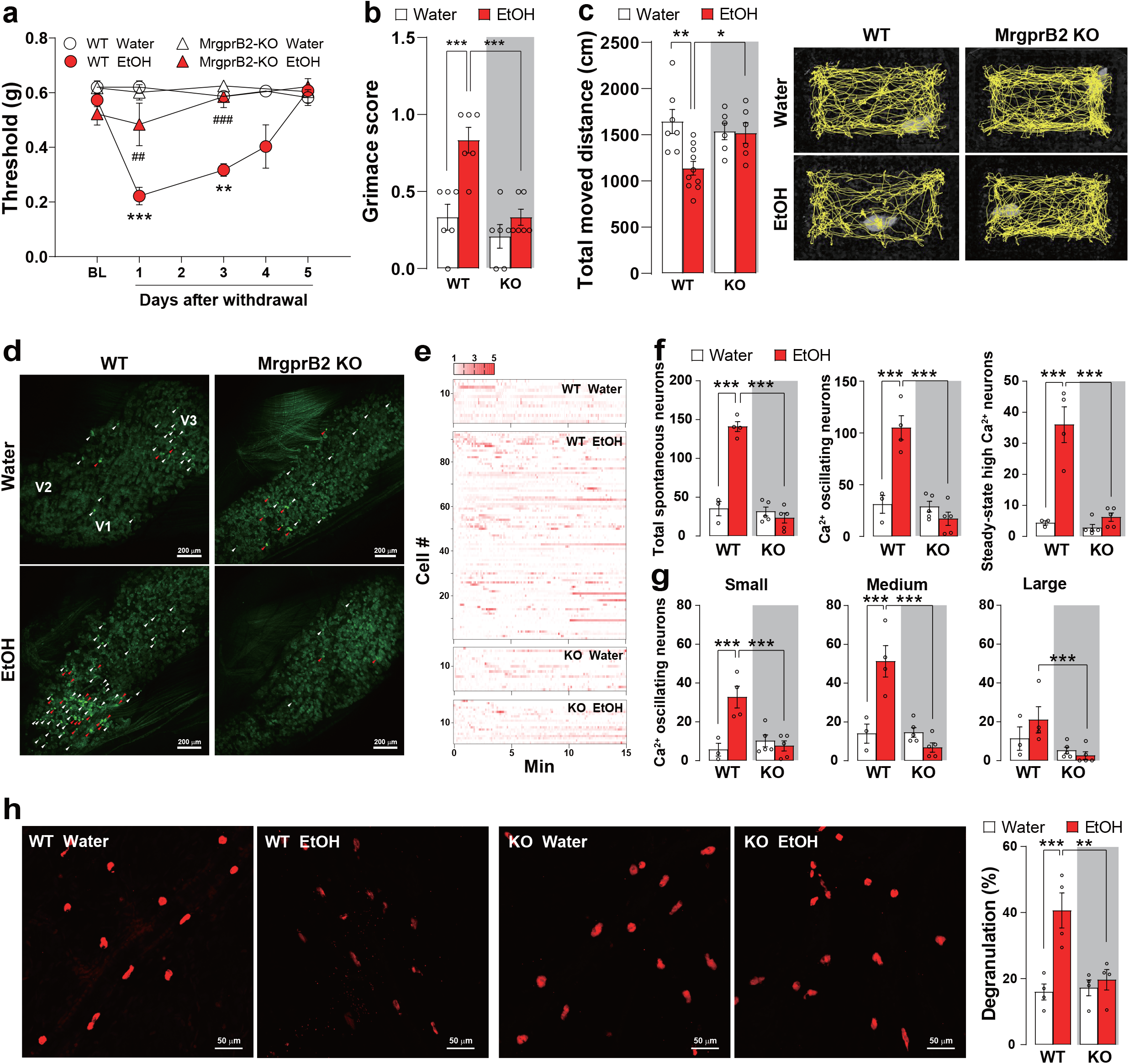
Mast cell degranulation via MrgprB2 contributes to alcohol withdrawal-induced headache and pain behaviors and sensitization of TG neurons. **a**, Facial mechanical withdrawal thresholds at various time points after alcohol withdrawal following 3 weeks of voluntary ethanol consumption vs. water only control in wild type (WT) or MrgprB2-deficient (KO) mice (n = 6 per group). **b**, Grimace score measured 24 hours after alcohol withdrawal (n = 6 per group). **c**, (*left*) Total distance moved in open field in 5 min measured 24 hours after alcohol withdrawal. (*right*) Representative traces of total moved distance in 5 mins (n = 7-10). **d**, Representative images of spontaneous activities in *in vivo* TG Pirt-GCaMP3 Ca^2+^ imaging after alcohol withdrawal. V1 (ophthalmic), V2 (maxillary), and V3 (mandibular) indicate location of neuronal cell bodies in TG image. Spontaneously Ca^2+^ oscillating neurons (white arrowheads). Spontaneously steady-state high Ca^2+^ activated neurons (red arrowheads). **e**, Representative heatmaps of spontaneously activated individual neurons. **f**, Number of total, oscillating, and steady-state high Ca^2+^ spontaneously activated neurons (n = 3-5). **g**, Number of Ca^2+^ oscillating neurons according to cell diameter of each group. Small-diameter TG neurons (<20 μ m); medium (20-25 μm); large (>25 μm). **h**, (*left*) Representative confocal fluorescent mast cell images of dura mater. Mast cells (red) were identified by avidin staining. (*right*) Percent degranulation of mast cells from each group (n = 4). TG: trigeminal ganglia. Error bars indicate S.E.M. *p < 0.05; **p < 0.01; ***p < 0.001, one-way ANOVA with Dunnett’s multiple comparison post-hoc test.

### Mast cell activation via MrgprB2 in dura mater involve in alcohol withdrawal-induced sensitization of TG neurons

To assess the activity of TG neurons in mice following ethanol withdrawal after 3 weeks of alcohol access, we monitored the activity of TG neurons in intact live animals using *in vivo* TG Pirt-GCaMP3 Ca^2+^ imaging. The total number (>141±6.34) of spontaneously activated neurons was dramatically increased in TG of alcohol withdrawal mice compared to water-fed controls (Fig. 1d-f). The spontaneously activated neurons included neurons exhibiting Ca^2+^ oscillations and neurons with steady-state high Ca^2+^, both of which were significantly increased in alcohol withdrawal mice (Fig. 1d-f). The group of small-diameter (<20 μm) and medium-diameter (20-25 μm) neurons from alcohol withdrawal mice showed more spontaneously activated neurons than water-fed controls (Fig. 1g and Extended Data Fig. 1g). The numbers of spontaneously activated TG neurons were significantly decreased in alcohol withdrawal of MrgprB2-deficient mice (Fig. 1d-g). Ethanol consumption is known to modulate mast cell activities, including increasing degranulation^19– 21^. Indeed, degranulated mast cells and the total number of mast cells were increased in dura mater of ethanol drinking mice (Fig. 1h and Extended Data Fig. 2a), and these increases were abolished in MrgprB2-deficient mice (Fig. 1h and Extended Data Fig. 2a). These data indicate that MrgprB2 is required for mast cell degranulation evoked by alcohol withdrawal, suggesting that mast cell activation via MrgprB2 results in development of alcohol withdrawal-induced headache and pain behaviors.

### CRF-activated MrgprB2 in dura mater induces headache behavior

Our findings indicate that degranulation of mast cells via MrgprB2 sensitizes TG neurons to evoke alcohol withdrawal-induced periorbital mechanical hypersensitivity and pain behaviors, but it is still unclear how alcohol withdrawal mediates activation of MrgprB2 leading to degranulation of mast cells. We sought to identify a putative activator of MrgprB2 after alcohol withdrawal. Alcohol consumption and withdrawal is known to activate the hypothalamic-pituitary-adrenal (HPA) axis^23–27^. We postulated that Corticotropin-releasing factor (CRF), a hormone active in regulating HPA axis function, might be involved in MrgprB2 activation and subsequent development of alcohol withdrawal-induced headache^28,29^. We found significantly increased CRF localization and expression to the dura mater of alcohol withdrawal mice compared to water-fed controls (Fig. 2a and Extended Data Fig. 2b). Since CRF, a secretagogue of mast cells, induces mast cell degranulation^30^, we hypothesized that CRF-induced activation of mast cells mediates alcohol withdrawal-induced headache behaviors. To confirm whether local increases of CRF in dura mater induce headache-like and pain behaviors, CRF was directly injected into dura mater. CRF injection significantly induced periorbital mechanical hypersensitivity from 1 to 24 hrs post-injection, which returned to baseline within 72 hrs (Fig. 2b). This was similar to the effect of IL-6, which is a well-known periorbital mechanical hypersensitivity inducer^31,32^. However, MrgprB2-deficient mice did not show periorbital hypersensitivity following dural CRF injection (Fig. 2b). Recent studies reported that MrgprB2 responds to various positively charged protein-secretagogues^15,33^, which led us to test whether CRF directly activates MrgprB2. We confirmed that CRF induced Ca^2+^ transients in HEK293 cells expressing MrgprB2 or the human ortholog MrgprX2 (Fig. 2c,d). We also applied CRF onto isolated mouse mast cells and cultured LAD2 human mast cells *in vitro*, and found that the CRF application also evoked Ca^2+^ transients in isolated mouse mast cells and LAD2 human mast cells (Fig. 2e,f). To rule out the possibility that CRF indirectly activated MrgprB2 in mast cells by engaging CRF-induced Ca^2+^ transients via CRF receptors, we added astressin, a CRF 1/2 receptor inhibitor, before CRF application. Astressin treatment did not block CRF-induced Ca^2+^ transients in isolated mouse mast cells or LAD2 human mast cells but did increase Ca^2+^ transients (Fig. 2g,h). Consistent with this result, astressin injection into dura mater enhanced the effect of CRF on periorbital mechanical hypersensitivity (Extended Data Fig. 2c). In addition, using *in vivo* TG Pirt-GCaMP3 Ca^2+^ imaging, we found a significant increase in the number of activated neurons following dural CRF injection (Fig. 2i). In *in vivo* dura mater imaging, direct CRF application onto dura mater induced vasodilation (Fig. 2j), a primary event occurring during migraine and headaches^7,34^. These results indicate that CRF, which is likely released from dural blood vessels, induces mast cell degranulation via MrgprB2 activation, and regulates development of alcohol withdrawal-induced headache and pain behaviors by sensitization of TG nerve in dura mater.

**Fig. 2.**
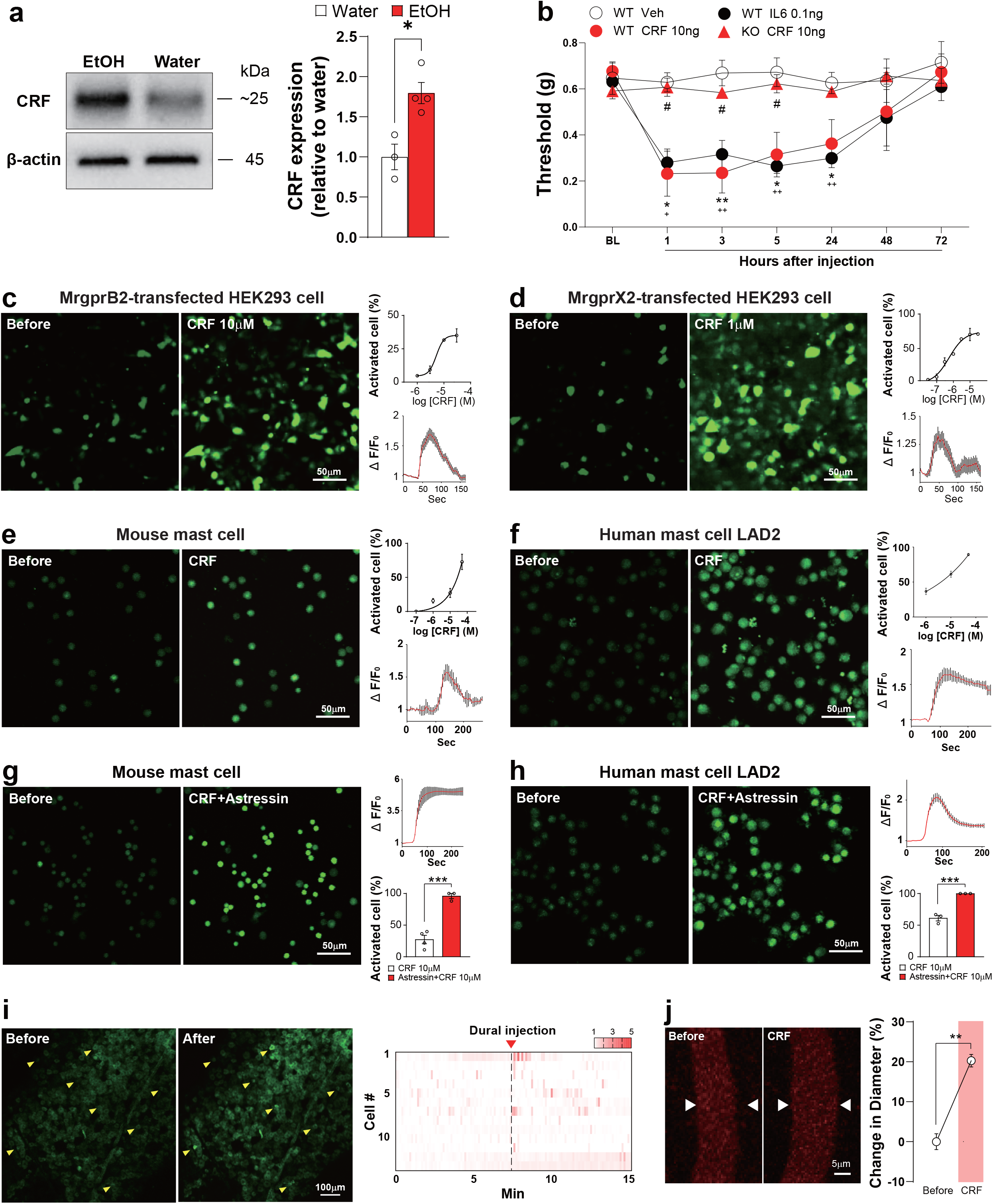
CRF-mediated mast cell activation via MrgprB2 induces headache behavior. **a**, (*left*) Representative western blot images and quantification (*right*) of CRF expression from dura mater after alcohol withdrawal (n = 15). **b**, Facial mechanical withdrawal thresholds after dural injection with IL-6 (0.1 ng) or CRF (10 ng) in wild type (WT) or MrgprB2-deficient (KO) mice (n = 4-7). *WT Veh. vs. WT CRF; ^+^WT Veh. vs. WT IL-6; ^#^WT CRF vs. KO CRF. **c** and **d**, (*left*) Representative Fluo-8 Ca^2+^ dye fluorescent images showing [Ca^2+^]_in_ increase in HEK293 cells expressing MrgprB2 or MrgprX2 before and after bath application of CRF (10 µM or 1 µM CRF). (*right, top*) Percentage of total cells activated by CRF bath application in MrgprB2- or MrgprX2-transfected HEK293 cells. (*right, bottom*) Average Ca^2+^ transient trace of cells activated by CRF from MrgprB2 (10 µM CRF)- or MrgprX2 (1 µM CRF)-transfected HEK293 cells (n = 3). **e** and **f**, (*left*) Representative Fluo-8 Ca^2+^ dye fluorescent images showing [Ca^2+^]_in_ increase by CRF (50 µM) bath application in isolated mouse mast cells (e) or LAD2 human mast cells (f). (*right, top*) Percentage of total cells activated by CRF bath application from isolated mouse mast cells (e) or LAD2 human mast cells (f). (*right, bottom*) Average Ca^2+^ transient trace of cells activated by CRF (50 µM) from isolated mouse mast cells (e) or LAD2 human mast cells (f) (n = 3). **g** and **h**, (*left*) Representative Fluo-8 Ca^2+^ dye fluorescent images showing [Ca^2+^]_in_ increase by CRF (10 µM) and astressin (10 µM) bath application in isolated mouse mast cells (g) or LAD2 human mast cells (h). (*right, top*) Average Ca^2+^ transient trace of cells activated by CRF (10 µM) and astressin (10 µM) from isolated mouse mast cells (g) or LAD2 human mast cells (h). (*right, bottom*) Percentage of total cells activated by CRF (10 µM) or CRF (10 µM) and astressin (10 µM) together from isolated mouse mast cells (g) or LAD2 human mast cells (h), (n = 3). **i**, (*left*) Representative TG images of spontaneous neuronal activities by direct CRF dural injection using *in vivo* TG Pirt-GCaMP3 Ca^2+^ imaging. (*right*) Heatmap showing [Ca^2+^]_in_ increase in TG neurons after direct CRF dural injection. Yellow arrowheads indicate activated TG neurons after CRF (10 ng) injection in dura. **j**, (*left*) Representative *in vivo* fluorescent dura blood vessel (red) images stained by Texas red dye using tail vein injection. (*right*) Percentage change in blood vessel diameter after CRF (20 µM) application in dura mater through cranial window. Dura blood vessel images were captured by confocal microscopy. CRF: corticotropin-releasing factor. TG: trigeminal ganglia. Error bars indicate S.E.M. *p < 0.05; **p < 0.01 by (a and j) two-tailed unpaired Student’s t test and (b) two-way ANOVA with Bonferroni’s multiple comparison post-hoc test.

### MrgprB2 contributes to alcohol withdrawal-induced sensitization of TG nerves

From studies identifying the pathophysiology of migraine headache, it is known that cutaneous allodynia is observed by non-noxious stimuli to periorbital and forehead skin areas during headache^35^. To confirm sensitization of TG nerve with various stimuli, including mechanical, thermal, and chemical stimuli, we applied von Frey filament (mechanical), hot water (thermal), or capsaicin (chemical) to each orofacial region innervated by ophthalmic (V1), maxillary (V2), or mandibular (V3) branches of TG nerves during *in vivo* TG Pirt-GCaMP3 Ca^2+^ imaging. Application of 0.4 g to the V1 region, but not to V2 or V3 regions, revealed increased activation of TG neurons in alcohol withdrawal mice (Fig. 3b and Extended Data Fig. 3a,h). This signal was due to increased activation of small-diameter to medium-diameter neurons (Fig. 3b). Mild hot water (40°C) or acetone (cold stimulus) applications also revealed greater numbers of activated TG neurons in V1 region but not in V2 or V3 regions (Fig. 3c and Extended Data Fig. 3C,J,E,L). The number of small to medium-diameter neurons activated in V1 region was increased in response to mild hot water (40°C) in alcohol withdrawal mice (Fig. 3c). Capsaicin injection into the V1 region but not in V2 or V3 regions of alcohol withdrawal mice resulted in an increase in activated TG neurons (Fig. 3d and Extended Data Fig. 3g,n). The sensitization of TG neurons in V1 region were also observed in alcohol withdrawal mice withdrawn from voluntary ethanol consumption after as long as 8 weeks (Extended Data Fig. 3p,s). Consistent with the results of headache and pain behaviors, MrgprB2-deficient mice after alcohol withdrawal did not exhibit an increase in activated TG neurons relative to control groups in response to all stimuli in all three branches (Fig. 3b-d). Collectively, these results suggest that alcohol withdrawal mice exhibit sensitization of TG nerve by mechanical, thermal, and chemical stimuli, and suggest that the mast cell-specific receptor MrgprB2 contributes to alcohol withdrawal-induced sensitization of TG nerves responsible for inducing headache and pain behaviors.

**Fig. 3.**
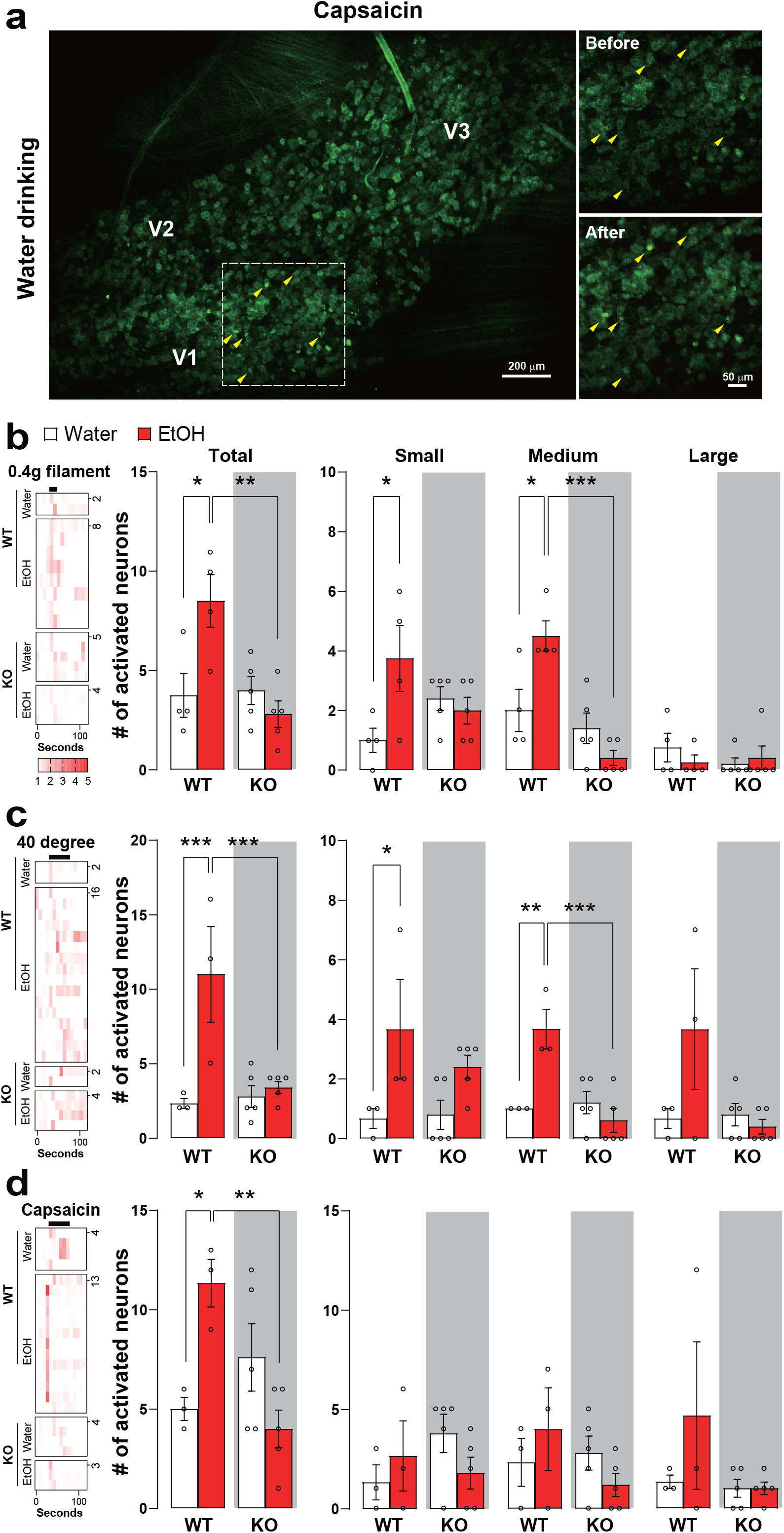
MrgprB2 deletion prevents sensitization (mechanical, thermal, and chemical) of TG neurons after alcohol withdrawal. **a**, Representative images of *in vivo* TG neurons activated by direct capsaicin (500 µM, 10 µL) injection into V1 region in water-fed control mice. Capsaicin was subcutaneously injected into the V1 branch (periorbital) region using an insulin syringe; *In vivo* TG Pirt-GCaMP3 Ca^2+^ imaging was performed after alcohol withdrawal. V1 (ophthalmic), V2 (maxillary), and V3 (mandibular) indicate location of neuron cell bodies in TG image. Yellow arrowheads indicate activated TG neurons after capsaicin injection into V1 region. **b-d**, Heatmaps and number of individual TG neurons activated by 0.4 g von Frey filament (b), mild hot water (40°C), or capsaicin (c and d) (n = 4-5). Number of activated neurons in response to various stimuli was plotted according to the size (small, medium, large) of each member of the pair. Small-diameter TG neurons (<20 µm); medium (20-25 µm); large (>25 µm). TG: trigeminal ganglia. Error bars indicate S.E.M. *p < 0.05; **p < 0.01; ***p < 0.001, one-way ANOVA with Dunnett’s multiple comparison post-hoc test.

### Alcohol withdrawal induces hypersensitivity of hindpaw and sensitization of DRG neurons

Consistent with previous studies^36–38^, during voluntary ethanol consumption for 3 to 8 weeks, mice developed hypersensitivity in hindpaw after withdrawal (Fig. 4a and Extended Data Fig. 4a), which lasted for 5 days following alcohol withdrawal. To examine whether sensitization of dorsal root ganglia (DRG) neurons occurs in mice following alcohol withdrawal, we also monitored neuronal activation by *in vivo* DRG Pirt-GCaMP3 Ca^2+^ imaging. Mild (100 g) or noxious (300 g) press to the hindpaw of alcohol withdrawal mice significantly increased the number of activated DRG neurons compared to water-fed controls (Fig. 4c-h). In response to these stimuli, small- and medium-diameter neurons were significantly more activated in DRG of alcohol withdrawal mice than water-fed controls (Fig. 4e,h). We also found an increase in activated neurons in the DRG of alcohol withdrawal mice in response to noxious heat (50°C) (Fig. 4i-k). The numbers of small- and medium-diameter neurons activated by noxious heat were significantly higher in alcohol withdrawal mice than in water-fed controls (Fig. 4k).

**Fig. 4.**
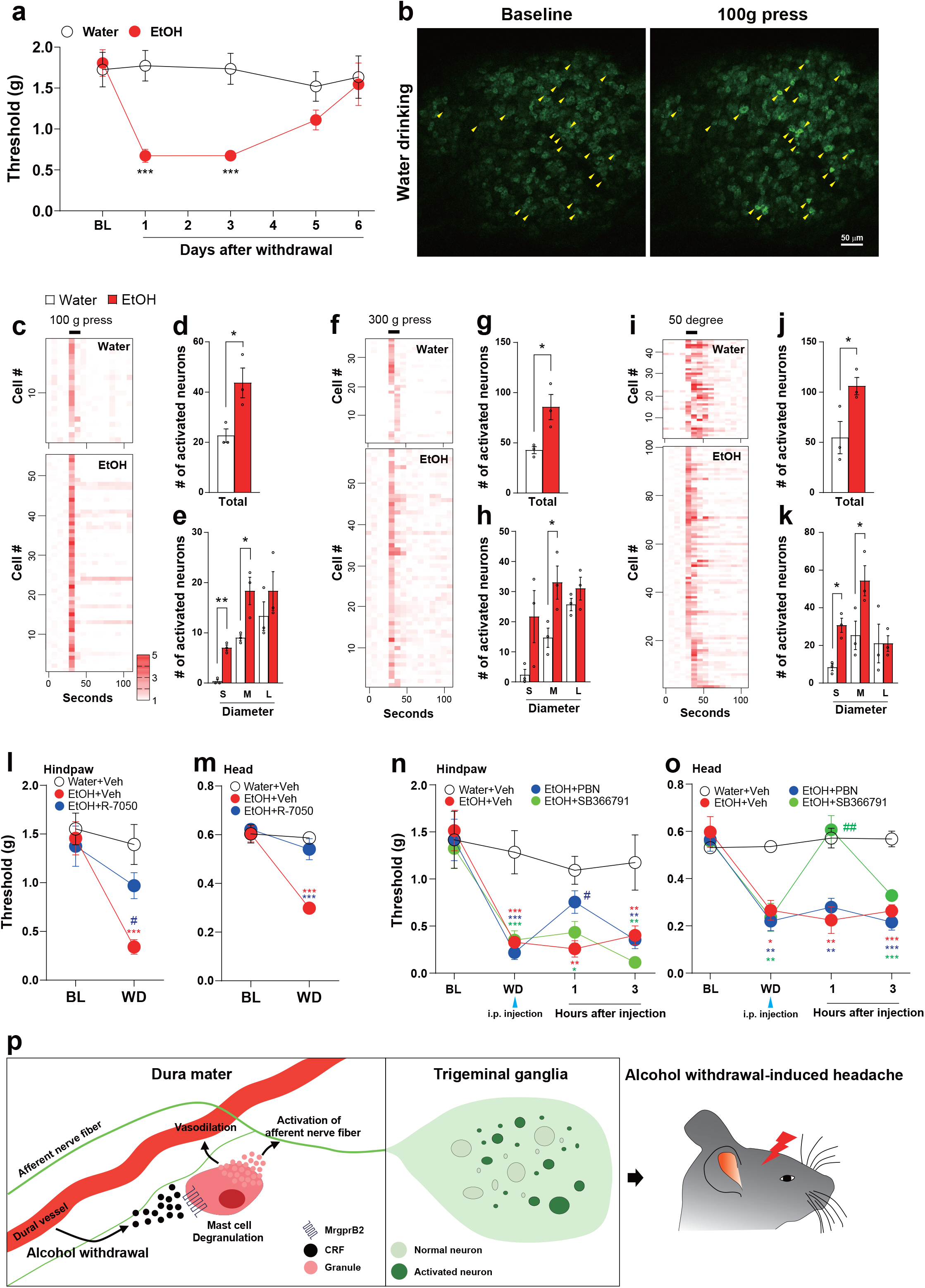
Head hypersensitivity and hindpaw hypersensitivity driven by alcohol withdrawal involve distinctive mechanisms. **a**, Hindpaw mechanical withdrawal thresholds at various time points after alcohol withdrawal following 3 weeks of voluntary ethanol consumption vs. water only controls (n = 9). **b**, Representative images of *in vivo* DRG Pirt-GCaMP3 Ca^2+^ imaging activated by 100 g hindpaw press in water-only control mice. Yellow arrowheads indicate DRG neurons activated by 100g hindpaw press. **c-k**, Heatmaps and number of individual neurons activated by 100 g press (c-e), 300 g press (f-h), or hot water (50°C, i-k) stimuli onto hindpaw (n = 3). S, Small-diameter DRG neurons (<20 µm); M, medium (20-25 µm); L, large (>25 µm). **l** and **m**, Hindpaw mechanical (l) and head withdrawal (m) thresholds tested by von Frey filament at 24 hours after alcohol withdrawal together with R-7050 (12 mg/kg) i.p. injection for 11 consecutive days (n = 4-6). *(red) Water+Veh. vs. EtOH+Veh.; *(blue) EtOH +Veh. vs. EtOH+R-7050; #(blue) EtOH+Veh. vs. EtOH+R-7050. **n** and **o**, Hindpaw mechanical (n) and head withdrawal (o) thresholds at 24 hours after alcohol withdrawal together with PBN (100 mg/kg) or SB366791 (1 mg/kg) i.p. injection (n = 4-5). *(red) Water+Veh. vs. EtOH+Veh.; *(blue) Water+Veh vs. EtOH+PBN; *(green) Water+Veh. vs. EtOH+SB366791; #(blue) EtOH +Veh. vs. EtOH+PBN; #(green) EtOH +Veh. vs. EtOH+SB366791. **p**, Schematic model of alcohol withdrawal-induced headache. Alcohol withdrawal-induced CRF release from blood vessels could induce mast cell degranulation in dura mater, which results in sensitization of primary sensory TG neurons and vasodilation. This pathway ultimately induces headache behavior. DRG: dorsal root ganglia. TNF-α receptor: tumor necrosis factor-α receptor. TRPV1: transient receptor potential channel V1. ROS: reactive oxygen species. CRF: corticotropin-releasing factor. TG: trigeminal ganglia. Error bars indicate S.E.M. *p < 0.05; **p < 0.01; ***p < 0.001, (a-k) two-tailed unpaired Student’s t test; (l-m) one-way ANOVA with Dunnett’s multiple comparison post-hoc test; (n and o) two-way ANOVA with Bonferroni’s multiple comparison post-hoc test.

### Alcohol withdrawal-induced headache behavior is mediated by TNF-α and TRPV1

To identify specific molecular signalling mechanisms that drive hypersensitivity in different anatomical locations as a result of alcohol withdrawal, we investigated whether inhibitor of tumour necrosis factor-α (TNF-α) receptor, which was increased by alcohol consumption^39^, would block alcohol withdrawal-induced mechanical allodynia. We found that alcohol withdrawal-induced mechanical allodynia in hindpaw or in the head was blocked by R-7050, a TNF-α receptor inhibitor (i.p. injection for 11 consecutive days) (Fig. 4l,m). A recent study suggested that chronic alcohol consumption induces nociceptor sensitization in hindpaw through an increase in reactive oxygen species (ROS)^37^, which led us to ask whether ROS production is a consequence of TNF-α release, and if ROS are involved in both hindpaw and head mechanical allodynia. We found that mechanical allodynia in hindpaw of alcohol withdrawal mice was reversed by PBN, a ROS scavenger, whereas mechanical allodynia in the head was not reversed (Fig. 4n,o). We also confirmed that the mechanical allodynia in the head caused by alcohol withdrawal was reversed by SB366791, a TRPV1 inhibitor (Fig. 4n,o). Activated mast cells can release TNF-α, which is a well-known potentiator of TRPV1 ion channel^40^, suggesting that in alcohol withdrawal mice TRPV1 activation is a component of a downstream signalling cascade of mast cell activation via MrgprB2. These mechanisms are independent pathways from CRF receptor signaling.

## Discussion

Clinical observations suggested that alcohol drinking causes headaches^41–44^, but the pathophysiological mechanism still remains unknown. Here, we show that withdrawal from alcohol drinking causes headache and pain behaviors, which are associated with sensitization of TG neurons. We also show that CRF directly activates MrgprB2, which mediates alcohol withdrawal-induced behavioral and cellular changes (Fig. 4p). Moreover, we identified different signaling mechanisms of alcohol withdrawal-induced hypersensitivity in the head compared with hypersensitivity in hindpaw. These results identify a mechanism of alcohol withdrawal-induced headache, and point to a therapeutic target for treating alcohol withdrawal-induced headache problems and addictive behaviors.

Mast cells in dura mater participate in inflammation and then sensitization of peripheral afferents, which are considered as a cascade of the development of migraine headache^12,45^. We here demonstrate that CRF activates mast cells via Mrgprb2, which causes headache behaviors and the TG nerve sensitization. These results indicate that CRF mediates alcohol withdrawal-induced physiological changes and mast cell degranulation that produces headache. Since alcohol withdrawal can induce changes in various hormones or peptides, there is a possibility that upregulation or downregulation of other hormones or peptides can affect alcohol withdrawal-induced behavioral and cellular changes via mast cell activation. Mast cells express CRF1/2 receptors. Unlike Mrgprb2, CRF receptors act as modulators and/or indirectly affect mast cell activity. It has been reported that CRF1 receptor enhances calcium transient and calcium signal and CRF2 receptor suppresses it in mast cells^46^. We here confirm that the inhibition of CRF1/2 receptors enhances CRF-induced mast cell activation. Because expression of CRF receptors is changed under various conditions^46^, further study is needed about the modulatory role of CRF receptors in mast cells of alcohol withdrawn mice. The withdrawal-induced physiological changes such as CRF elevation also occur in the use of other addictive substances, including heroin and other opioids^47^. Thus, CRF-MrgprB2 axis-induced mast cell activation might be a significant signaling pathway in the development of headaches induced by addictive substances.

Pain and alcohol dependence have a close relationship. Acute pain-inhibitory effect by alcohol consumption and withdrawal from alcohol drive to motivate alcohol drinking, which induces severe alcohol dependence^2,3^. Indeed, patients receiving treatment for alcohol use disorder exhibit more significant pain responses during the early stages of alcohol abstinence. In adults with chronic pain, their pain intensity is correlated with increased alcohol consumption^3^. Therefore, pain relief, especially from headaches could be a way to lower severe alcohol dependence^5^.

Alcohol abuse or addiction is a significant public health problem, especially during the COVID-19 pandemic because the pandemic is leading people to drink alcohol a lot more than before. Indeed, the pandemic increases vulnerability to the development of alcohol addictive behaviors^1^. Moreover, pains such as headaches from alcohol withdrawal disrupt rehabilitation from alcohol addiction and/or abuse^3,5^. We here demonstrate that the blockade of mast cell-specific MrgprB2 in mast cells attenuates alcohol withdrawal-induced headache behaviors. Therefore, our results provide CRF/MrgprB2 as new therapeutic targets for treating alcohol withdrawal-induced headaches and even for alcohol addiction and/or abuse.

## Methods

### Animals

The C57BL/6 mice used in experiments were obtained from The Jackson Laboratory; Pirt-GCaMP3 mice were generated and used as described previously^48,49^. MrgprB2-deficient (KO) mice on a C57BL/6 background were purchased from The Centre for Phenogenomics (Ontario, Canada). All mice spent at least one week in group housing (4-5 mice per cage, 23°C, regular light/dark cycle), and then were split into single housing one week before the experiment. Mice used for experiments were 8-16 weeks old. All experiments were performed in accordance with a protocol approved by the University of Texas Health at San Antonio (UTHSA) Animal Care and Use Committee (IACUC).

### Peptides and drugs

CRF peptide was purchased from Phoenix Pharmaceuticals (Burlingame, CA, USA). R-7050, PBN, SB366791, and astressin were purchase from Tocris Bioscience/ Bio-Techne Corporation (Minneapolis, MN, USA).

### Alcohol withdrawal model

The voluntary two-bottle choice ethanol drinking paradigm was used for ethanol self-administration^36^. Mice received ad libitum 24 hours access for 3 weeks or for 8 weeks to two bottles: one bottle containing water and another bottle containing ethanol. The location of the bottles on the cage was exchanged every other day. The alcohol bottle contained 3% ethanol (days 1-2), 6% ethanol (days 3-4), and 10% ethanol thereafter.

**Behavior tests (mechanical, grimace, open field). Hindpaw or facial mechanical von Frey test** was performed according to previously published methods^32,36^. Mice were familiarized and habituated to the experimenter’s smell, hand touch, and eye contact for at least for 3 days, and then were acclimated in a plexiglass chamber with 4 oz paper cups for 2 h/d for 3 days. After acclimation, mice were subjected to baseline testing of cutaneous facial (periorbital region) or hindpaw sensitivity touched by von Frey filament for approximately 5 days. Baseline of the facial von Frey test was defined as a withdrawal threshold of approximately 0.5 to 0.7 g. The thresholds were determined using the Dixon “up-and-down” method^50^. **Grimace pain behavior** was recorded in five characterized facial areas (orbital, nose, cheek, ears, and whiskers) on a scale of 0 to 2 (0 = not present, 1 = somewhat present, 2 = clearly present) as previously published^51^. **The open field test** was performed in a new cage for 5 min. The mouse’s movement was recorded using a video camera, and the movement was analysed using ImageJ (NIH) with animal tracker (plugin)^52^.

### DRG and TG exposure surgery for in vivo DRG and TG Pirt-GCaMP3 Ca^2+^ imaging

**DRG exposure surgery** was performed as previously described^48^. Mice were anesthetized by i.p. injection of Ketamine/Xylazine (approximately 80/10 mg/kg) (MilliporeSigma, St Louis, MO, USA), their backs were shaved and ophthalmic ointment (Lacri-lube; Allergen Pharmaceuticals) was applied to the eyes. The surface aspect of the L5 DRG transverse process was removed to expose underlying DRG. Bleeding from the bone was stopped using styptic cotton. **For TG exposure surgery**, we first surgically exposed the right side dorsolateral skull by removing skin and muscle. The dorsolateral skull (parietal bone between right eye and ear) was removed using a dental drill (Buffalo Dental Manufacturing, Syosset, NY, USA) to make a cranial window hole (∼10×10 mm). The TG was then exposed where it is located under the brain by aspirating overlying cortical tissue through a cranial window in the dorsolateral skull. The animal was then laid on its abdomen on the stage under a Zeiss LSM 800 confocal microscope (Carl Zeiss). The animal was restrained using a mouse tooth holder to minimize movements from breathing and heartbeats. During the surgery, the body temperature of the mouse was maintained on a heating pad at 37°C ± 0.5°C as monitored by rectal probe.

### In vivo Pirt-GCaMP3 Ca^2+^ imaging in DRG and TG

*In vivo* Pirt-GCaMP3 Ca^2+^ imaging in live mice was performed for 1-5 hours immediately after exposure surgery as previously described^48,49^. After the exposure surgery, mice were laid abdomen-down on a custom-designed platform under the microscope objective. For ***in vivo* DRG Pirt-GCaMP3 Ca**^**2+**^ **imaging**, the spinal column was stabilized using two clamps on vertebra bone above and below the DRG being imaged. Live images were acquired at ten frames per cycle in frame-scan mode per ∼4.5 to 8.79 s/frame, at ranging from 0 to 90 μm, using a 10 × 0.4 NA dry objective at 512 × 512 pixel or higher resolution with solid diode lasers tuned at 488 nm wavelength and emission at 500-550 nm. An average of 1,825±71 neurons per DRG (∼10-15% of total DRG neurons) was imaged. For ***in vivo* TG Pirt-GCaMP3 Ca**^**2+**^ **imaging**, the animal’s head was fixed by a custom-designed head holder. During the imaging session, body temperature was maintained at 37°C ± 0.5°C on a heating pad and monitored by rectal probe. Anaesthesia was maintained with 1-2% isoflurane using a gas vaporizer, and pure oxygen was delivered through a nosecone. Live images were acquired at ten frames per cycle in frame-scan mode per ∼4.5 to 8.79 s/frame, at ranging from 0 to 90 μm, using a 5 × 0.25 NA dry objective at 512 × 512 pixel or higher resolution with solid diode lasers tuned at 488 nm wavelength and emission at 500-550 nm. An average of 2,867±87 neurons per TG (∼10% of total TG neurons^53,54^) was imaged and small regions of TG neurons were imaged at faster speed >40Hz. von Frey filaments (0.4 g, and 2.0 g) were applied to the face or hindpaw of exposed TG branches or DRG side. 100 g and 300 g press were applied to the whole palm of hindpaw using rodent pincher (Bioseb, U.S.A.). Whole hindpaw or different TG branches at animal’s face was applied by 40 °C, 50 °C, or 60 °C water and acetone was applied by pipette to the hindpaw or the different TG branches. Capsaicin (500 μm, 10 μl) or KCl (500 mM, 10 μl) was cutaneously injected into the different TG branches using a 0.5-ml insulin syringe with a 28-gauge needle.

### In vivo Pirt-GCaMP3 Ca^2+^ imaging data analysis

For imaging data analysis, raw image stacks were collected, deconvoluted, and imported into ImageJ (NIH). Optical planes from sequential time points were re-aligned and motion corrected using the stackreg rigid-body cross-correlation-based image alignment plugin. Ca^2+^ signal amplitudes were expressed as F_t_/F_0_ as a function of time. F_0_ was defined as the average fluorescence intensity during the first two to six frames of each imaging experiment. All responding cells were verified by visual examination using the raw imaging data. Spontaneous activity from primary sensory neurons has two distinct events, Ca^2+^ oscillating neurons and steady-state high Ca^2+^ neurons without any stimuli. Ca^2+^ oscillating neurons are defined as Ca^2+^ intensity level changes or fluctuations over time. Steady-state high Ca^2+^ neurons are defined as Ca^2+^ intensity level highly increased from baseline maintained at the same or similar level over time (no fluctuation).

### Mouse dural injection

Mouse dura injection was performed as previously described^31^. Mice were anesthetized under isoflurane briefly, and drugs or compounds were injected in a volume of 5 µl via a modified internal cannula (P1 Technologies, Roanoke, VA, USA). The length of injection needle was adjusted to 0.5 to 0.65 mm.

### In vivo blood vessel imaging in dura mater

Mice were anesthetized by i.p. injection of Ketamine/Xylazine (80/10 mg/kg) (MilliporeSigma, St Louis, MO, USA) and ophthalmic ointment was applied to eyes. The scalp was shaved and was sterilized using 70% EtOH. For cranial window formation, a ∼3×3 mm round area of right parietal skull was carefully removed. Using dental cement (Lang Dental Manufacturing, Wheeling, IL, USA), a crown was made around the cranial window, and the brain was covered with PBS. Texas-Red conjugated 2k Dalton dextran (Nanocs) was injected into the tail vein (100 μl of 1 mg/ml in saline) to visualize blood vessels in dura mater. Anesthetized mice were imaged with a single photon confocal microscope (Carl Zeiss) using the 40X water immersion with 1.0 NA objective. The vessel diameter was measured using ImageJ software.

### Immunofluorescence imaging in dura mater

Animals were anesthetized with Ketamine/Xylazine (80/10 mg/kg) (MilliporeSigma, St Louis, MO, USA) before cardiac perfusion fixation in 4% paraformaldehyde. After fixation, the dura mater was dissected and postfixed for 24 hours in 4% paraformaldehyde. The collected tissues were incubated with primary antibodies: CRF (Santa Cruz), βIII-tubulin (Abcam), and Rhodamine Avidin D (Vector laboratories) for 24 hours. Tissues were then incubated with the fluorescent-dye labelled polyclonal secondary antibodies; Alexa-Fluor 488 or 647 (1:500; Invitrogen), and slides were cover-slipped using mounting medium (Invitrogen).

### Mast cell culture and imaging

Peritoneal mast cells were isolated from adult male and female mice as previously described^15^. Briefly, 2 × 5 mL of ice-cold mast cell dissociation medium (MCDM; HBSS with 3% FBS and 10 mM HEPES, pH7.2) was injected into the peritoneal cavity, and the abdomen was massaged for 60s; peritoneal fluid was collected, and centrifuged at 200 rcf for 5 min at room temperature. The pellets were resuspended in 2 mL MCDM, and then layered over 4 mL of an isotonic 70% Percoll solution (MilliporeSigma, St Louis, MO, USA) and centrifuged at 500 rcf for 20 min at 4°C. Purity of isolated mast cells was >95%, as determined from avidin staining. The mast cells were resuspended in DMEM with 10% FBS, 100 U/mL penicillin, 50 mg/mL streptomycin, and 25 ng/mL recombinant mouse stem cell factor (mSCF; Peprotech, Cranbury, NJ, USA), and plated onto glass coverslips coated with 30 mg/mL fibronectin (MilliporeSigma, St Louis, MO, USA). LAD2 human mast cells (Laboratory of Allergic Diseases 2), provided by Professor Dr. Xinzhong Dong from Johns Hopkins University (MD, USA), were cultured in StemPro-34 SFM medium (Life Technologies, Carlsbad, CA, USA) supplemented with 2 mM L-glutamine, 100 U/ml penicillin, 50 mg/ml streptomycin, and 100 ng/ml recombinant human stem cell factor (Peprotech, Cranbury, NJ, USA), and were maintained at 37°C, 5% CO_2_. Cell culture medium was hemi-depleted every week and replaced with fresh medium. For Ca^2+^ imaging, the cell suspensions were seeded onto glass coverslips as above at a density of 5,000 cells, and incubated for 2 hours (37°C, 5% CO_2_). The cells were loaded with Fluo-8 Ca^2+^ dye (5 µM) for 30 minutes and then imaged in Ca^2+^ imaging buffer (CIB; 125 mM NaCl, 3 mM KCl, 2.5 mM CaCl_2_, 0.6 mM MgCl_2_, 10 mM HEPES, 20 mM glucose, 1.2 mM NaHCO3, 20 mM sucrose, adjusted to pH 7.4 with NaOH) using a confocal (Carl Zeiss, Oberkochen, Germany) or widefield fluorescence microscope (Carl Zeiss) system.

### Ca^2+^ imaging in HEK293 Cells

HEK293 cells were cultured in DMEM-FBS media and maintained in a 5% CO_2_ incubator at 37°C. Prior to transfection, cells were trypsinized and plated onto poly-D-lysine-coated coverslip in 35 mm culture dishes. Following overnight incubation, the cells were transiently transfected with lipofectamine 2000 (Invitrogen), using a total of 2 µg cDNA per dish (35 mm). Plasmid DNAs encoding the human MrgprX2 (pcDNA3.1) and mouse MrgprB2 (pcDNA3.1), provided by Professor Dr. Xinzhong Dong from Johns Hopkins University (MD, USA), were co-transfected with Gα15 (pcDNA3.1) at a ratio of 9:1. A surrogate expression marker, tdTomato (60 ng), was co-transfected along with other plasmids. Forty-eight hours following transfection, cells were loaded with AM esters of the Ca^2+^ indicators, Fluo-8 (1 µM; Abcam, Cambridge, MA, USA) along with Pluronic F-127 (0.04%; Life Technologies, Carlsbad, CA, USA) for 30–45 min at 37°C in the dark, in standard extracellular solution (SES) containing 140 mM NaCl, 4 mM KCl, 2 mM CaCl_2_, 1 mM MgCl_2_, 10 mM HEPES, 10 mM D-glucose, pH 7.40. Cells were viewed on an upright Zeiss Examiner/A1 microscope fitted with a 40× water-immersion objective (0.75-NA, 2.1-mm free working distance, Carl Zeiss) and with an Axiocam 705 color camera (Carl Zeiss). Fluorescence images were taken alternately every 5 s using the Zeiss Zen Blue software module. Cells were imaged in SES at room temperature; drugs were bath applied into the chamber following 10 cycles of baseline imaging, and responses were monitored for an additional 50 cycles. The percentage of cells responding to the CRF among total tdTomato-expressing cells was calculated to quantify the CRF response.

### Western blot for CRF of dura mater

Tissue lysates were prepared and were analyzed with western blot as previously described ^55,56^. Briefly, the collected dura mater was homogenized and centrifuged for 30 min at 12,000 rpm. The protein samples were separated by SDS-PAGE and then transferred to a PVDF membrane (Amersham, Buckinghamshire, UK), which was immunoblotted with anti-CRF (Santa Cruz, 1:500) and anti-β-actin (Cell Signaling, 1:5000)^57^. Each expression was quantitated using ImageJ (NIH). Protein expression was normalized to β-actin in the same sample.

### Statistical analysis

Group data were expressed as mean ± S.E.M. Two-tailed unpaired Student’s t test, one-way ANOVA tests, and two-way ANOVA tests were used to determine significance in statistical comparisons, and differences were considered significant at p < 0.05.

## Acknowledgements

We thank Gregory Dussor and members of the Gregory Dussor laboratory for advice on the facial von Frey test and dura injection. This work was supported by National Institutes of Health grants (NIDCR, DE026677 to Y.S.K.), University of Texas Health Science Center at San Antonio (UTHSA) startup fund (Y.S.K.), and a Rising STAR Award from University of Texas System (Y.S.K.).

## Author contributions

H.S. and Y.S.K contributed to study design with assistance from A.A. H.S., J.S., A.A., and Y.S.K. contributed to data interpretation and manuscript revision. H.S. conceived the project and performed all experiments except where noted, and drafted the paper. Z.Y. performed HEK293 cell work. H.S. and J.S. maintained, set up mating, took care of mice, and performed genotyping. H.I. and R.G. assisted with von Frey test work. Y.S.K. supervised all aspects of the project and wrote the paper.

## EXTENDED DATA FIGURE LEGENDS

**Extended Data Fig. 1. Voluntary ethanol consumption over 3 weeks results in development of ethanol preference, and headache and pain behaviors without changes in body weight or food intake. a**, Ethanol preference developed with voluntary ethanol consumption over 3 weeks. Ethanol preference (%) was determined as (EtOH intake/total fluid intake) Χ 100) and was compared with water preference (%) determined as (water intake 2^nd^ bottle/total fluid intake) Χ 100). **b**, Absolute amount of ethanol intake per a day was measured (EtOH intake (g)/body weight (kg)/7 days). **c**, Food intake was measured for every week. **d**, Animal’s percentage body weight gain determined as (daily body weight/baseline body weight) X 100. **e**, Representative images of animal’s face grimace expression. **f**, Average velocity (cm/sec) was calculated in the open field test. **g**, Numbers of steady-state high Ca^2+^ activated neurons were counted according to the size of each member of the pair. Small-diameter TG neurons (<20 µm); medium (20-25 µm); large (>25 µm). **h-k**, Parameters of ethanol intake by MrgprB2-deficient mice were measured as in (a-d): percentage of ethanol preference (h), absolute ethanol intake amount (i), food intake (j), and body weight gain (k). TG: trigeminal ganglia. Error bars indicate S.E.M. *p < 0.05; **p < 0.01; ***p < 0.001 by (a and h) two-tailed unpaired Student’s t test and (b, f, g, and i) one-way ANOVA with Dunnett’s multiple comparison post-hoc test.

**Extended Data Fig. 2. Mast cell numbers of wild-type and MrgprB2-deficient mice in dura mater after alcohol withdrawal and the effect of astressin, CRF1/2 receptor inhibitor on CRF-induced facial mechanical withdrawal thresholds. a**, (*left*) Representative confocal microscopic images of mast cells in dura mater. Mast cells were stained by avidin (white). (*right*) Mast cell numbers were counted 24 hours after alcohol withdrawal using image J software. **b**, (*left*) Representative confocal fluorescent CRF images in dura mater. (*right*) Fluorescence value of CRF immunohistochemical staining 24 hours after alcohol withdrawal (n = 4). **c** Facial mechanical withdrawal thresholds after dural injection with CRF (2 ng) or CRF (2 ng) + astressin (4 ng) (n = 5). Error bars indicate S.E.M. *p < 0.05; **p < 0.01; ***p < 0.001 by (a) one-way ANOVA with Dunnett’s multiple comparison post-hoc test and (b and c) two-tailed unpaired Student’s t test.

**Extended Data Fig. 3. Various stimuli in V2 and V3 branch regions of ethanol-withdrawn mice did not produce an increased activation of TG neurons, but stimulation of the V1 region after alcohol withdrawal following 8 weeks of ethanol consumption produced strong sensitization of TG neurons**.

Total number of activated TG neurons in response to 0.4 g and 2 g von Frey filament (mechanical), 40°C, 60°C, and acetone (thermal), KCl and capsaicin (chemical) in branch regions after alcohol withdrawal following 3 weeks of ethanol consumption: **a-g**, V2 branch region; **h-n**, V3 branch region. **o**, Facial mechanical withdrawal thresholds at various time points after alcohol withdrawal following 8 weeks voluntary ethanol consumption or water only (n = 7 for each group). **p-s**, Total number of activated TG neurons in response to 0.4 g filament (mechanical), 40°C and acetone (thermal), and capsaicin (chemical) onto the V1 region in mice after alcohol withdrawal following 8 weeks of ethanol consumption. Small-diameter TG neurons (<20 µm); medium (20-25 µm); large (>25 µm). TG: trigeminal ganglia. Error bars indicate S.E.M. *p < 0.05; **p < 0.01; ***p < 0.001 by two-tailed unpaired Student’s t test.

**Extended Data Fig. 4. Alcohol withdrawal following 8 weeks of ethanol consumption causes strong hypersensitivity (mechanical and thermal) in hindpaw and sensitization of DRG neurons. a**, Hindpaw mechanical withdraw thresholds plotted using von Frey filament at 1, 3, 5, 6 days after alcohol withdrawal. **b-d**, Numbers of individual TG neurons activated by 100 g press and 300 g press (mechanical), and hot water (50°C) (thermal). S, Small-diameter DRG neurons (<20 µm); M, medium (20-25 µm); L, large (>25 µm). DRG: dorsal root ganglia. Error bars indicate S.E.M. *p < 0.05; **p < 0.01; ***p < 0.001 by two-tailed unpaired Student’s t test.

## Supplementary Movies

**Movie 1-4, related to Fig. 1**. Representative imaging of *in vivo* whole TG neurons from ethanol or water drinking in wild type (WT) or MrgprB2-deficient (KO) mice. Spontaneous activity was imaged (no stimuli were applied). Movie 1: Spontaneous activity, WT-water drinking, Movie 2: Spontaneous activity, WT-ethanol drinking, Movie 3: Spontaneous activity, MrgprB2KO-water drinking, Movie 4: Spontaneous activity, MrgprB2KO-ethanol drinking.

**Movie 5-8, related to Fig. 3**. Representative imaging of *in vivo* whole TG neurons from ethanol or water drinking in WT or MrgprB2-deficient (KO) mice. Mild hot thermal stimulus (40 °C) was applied to V1 cutaneous skin area and was indicated in all of the movies. Movie 5: 40 °C, WT-water drinking, Movie 6: 40 °C, WT-ethanol drinking, Movie 7: 40 °C, MrgprB2KO-water drinking, Movie 8: 40 °C, MrgprB2KO-ethanol drinking.

**Movie 9 and 10, related to Fig. 4**. Representative imaging of *in vivo* whole DRG neurons from ethanol or water drinking mice. Hot thermal stimulus (50 °C) was indicated in all of the movies. Movie 5: 50 °C, WT-water drinking, Movie 6: 50 °C, WT-ethanol drinking,

