## Supplementary figures and images for "Mast cell-specific receptor/corticotropin-releasing factor axis regulates alcohol withdrawal-associated headache"

### Extended Data Figure 1

#### Extended Data Figure 1

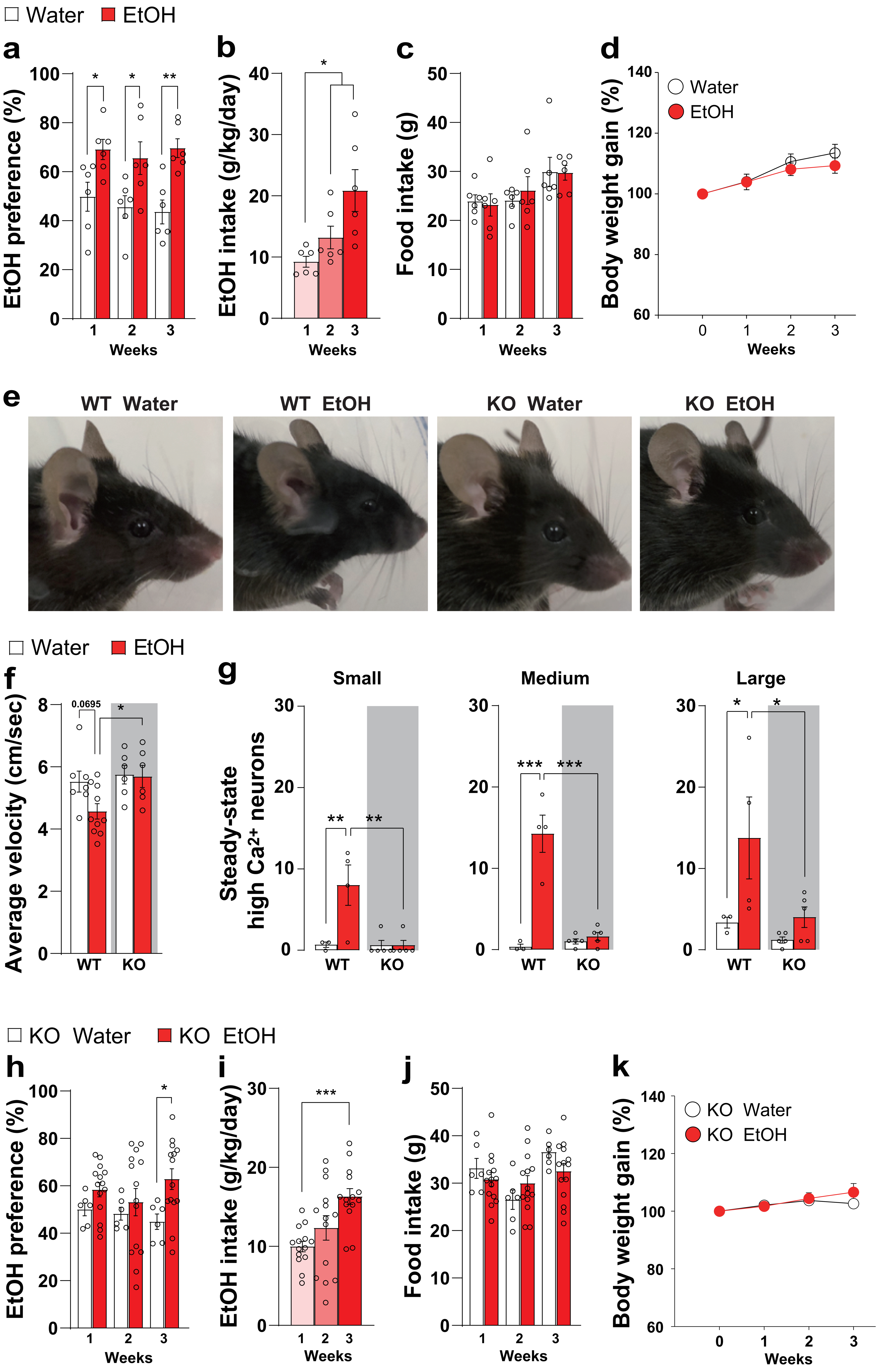

### Extended Data Figure 2

#### Extended Data Figure 2

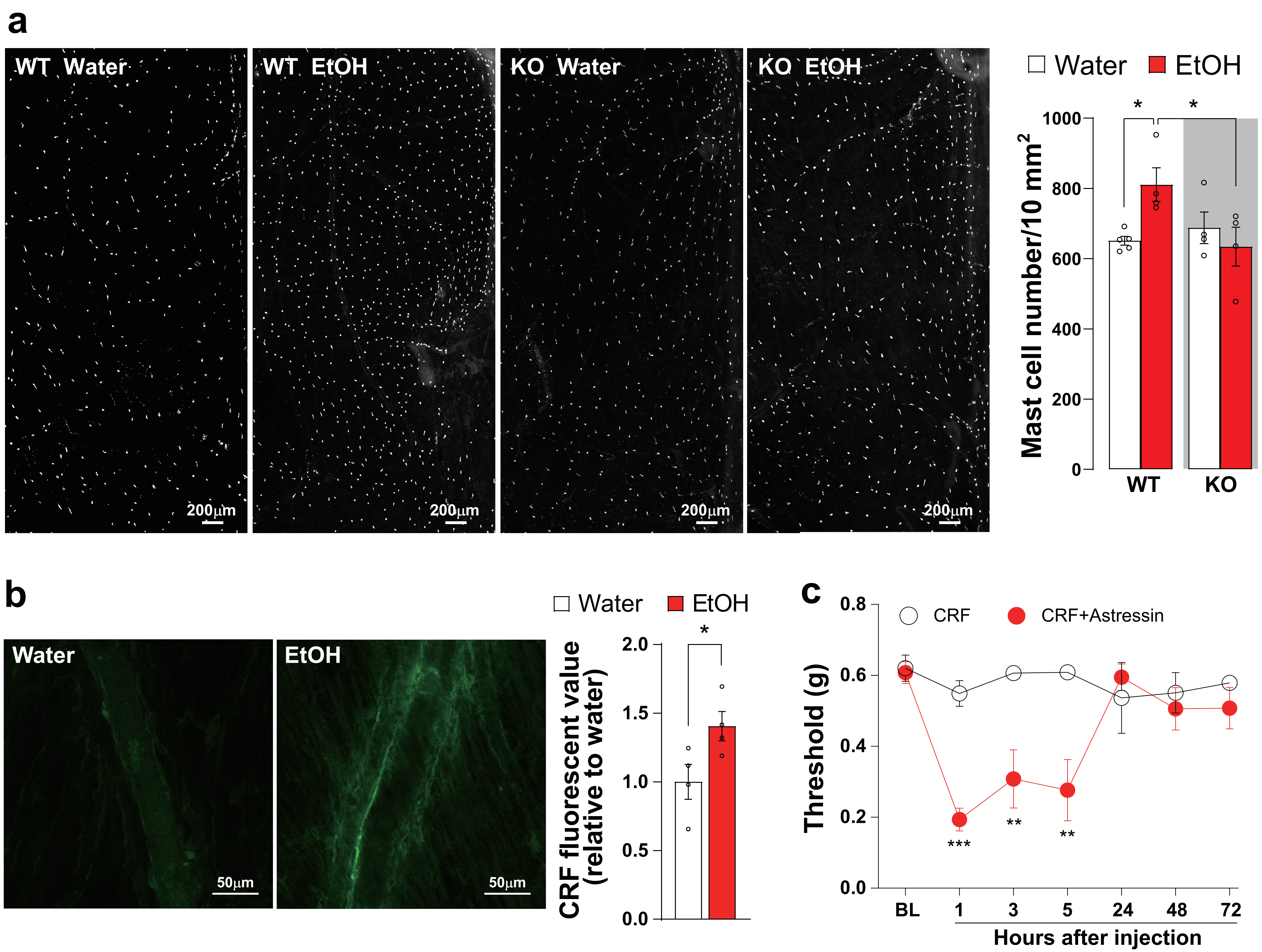

### Extended Data Figure 3

#### Extended Data Figure 3

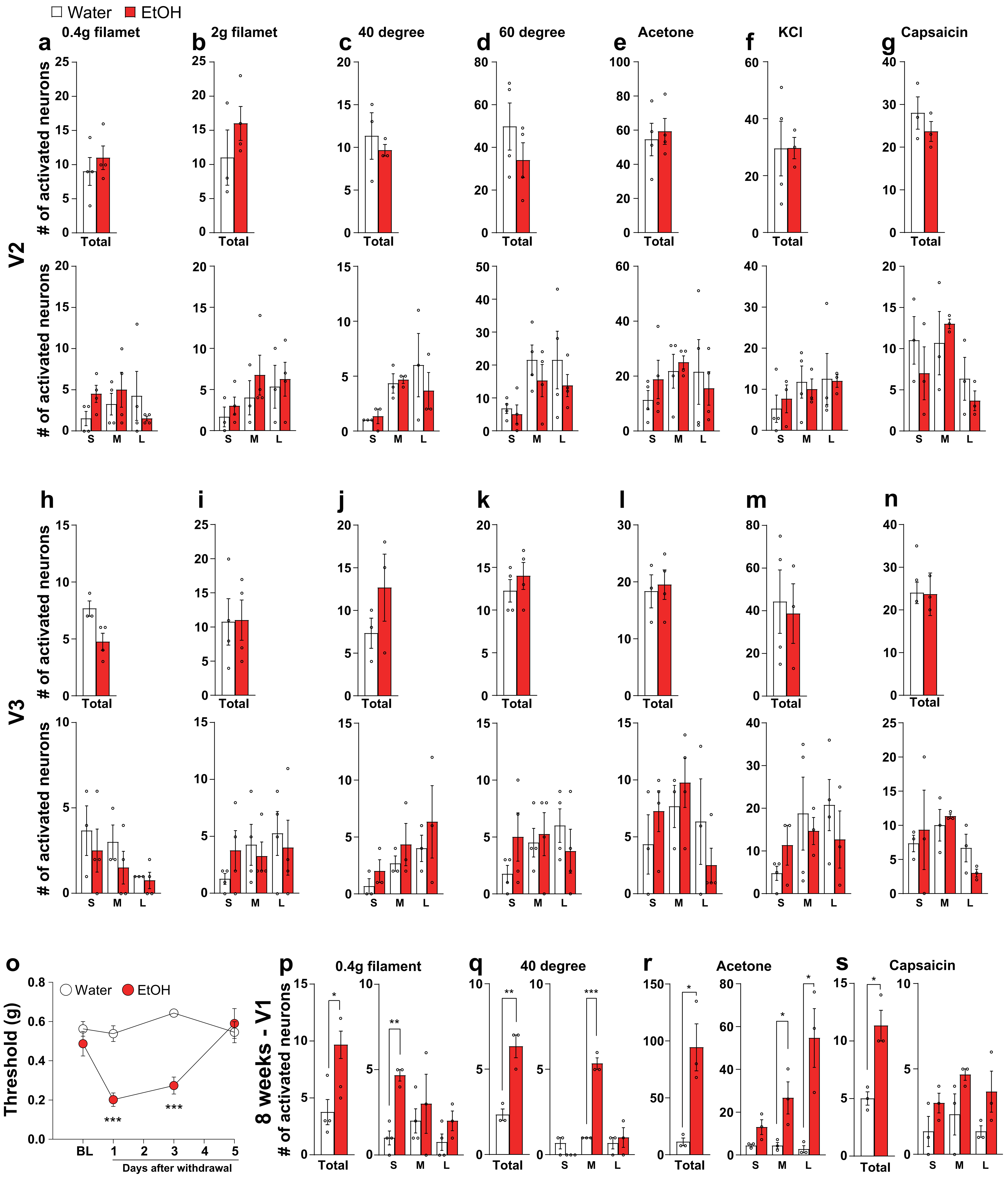

### Extended Data Figure 4

#### Extended Data Figure 4

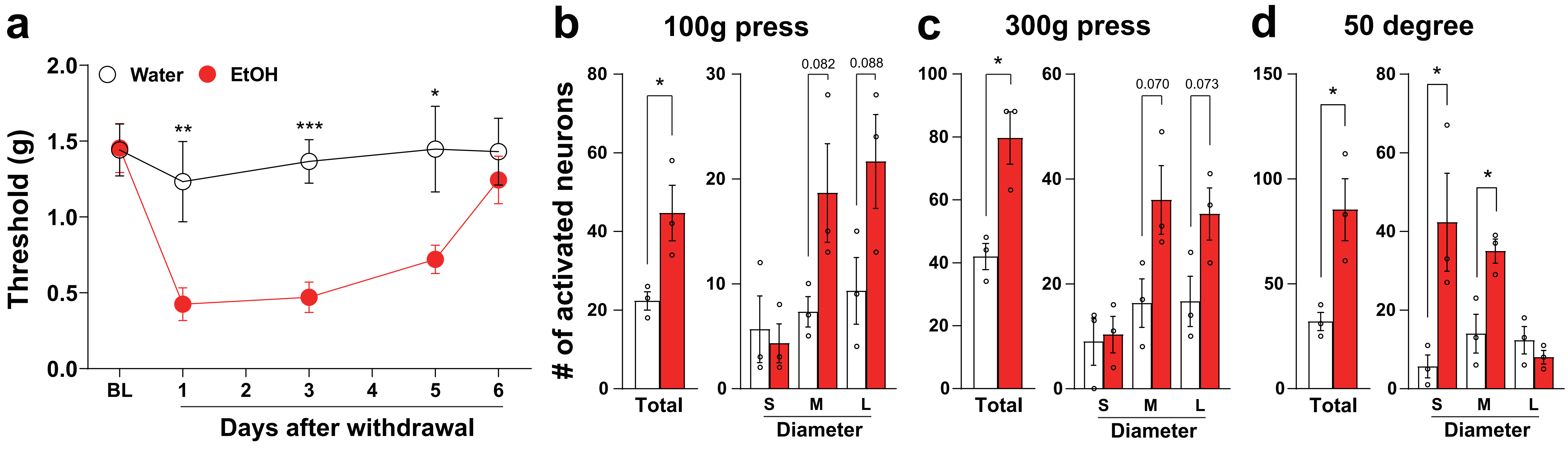
